# Exploration of polyyne biosynthetic gene cluster diversity in bacteria leads to the discovery of the *Pseudomonas* polyyne protegencin

**DOI:** 10.1101/2021.03.05.433886

**Authors:** Alex J. Mullins, Gordon Webster, Hak Joong Kim, Jinlian Zhao, Yoana D. Petrova, Christina E. Ramming, Matthew Jenner, James A. H. Murray, Thomas R. Connor, Christian Hertweck, Gregory L. Challis, Eshwar Mahenthiralingam

**Affiliations:** Microbiomes, Microbes and Informatics Group, Organisms and Environment Division, School of Biosciences, Cardiff University, Cardiff, CF10 3AX, UK; Department of Biomolecular Chemistry, Leibniz Institute for Natural Product Research and Infection Biology, Hans Knöll Institute, Beutenbergstrasse 11a, 07745, Jena, Germany; Department of Chemistry, University of Warwick, Coventry, CV4 7AL, UK; Warwick Integrative Synthetic Biology Centre, University of Warwick, Coventry CV4 7AL, UK; Molecular Biosciences Division, School of Biosciences, Cardiff University, Cardiff, CF10 3AX, UK; Faculty of Biological Sciences, Friedrich Schiller University Jena, 07743 Jena, Germany; Department of Biochemistry and Molecular Biology, Biomedicine Discovery Institute, Monash University, Clayton, VIC 3800, Australia

## Abstract

Natural products that possess alkyne or polyyne moieties have been isolated from a variety of biological sources. In bacteria their biosynthesis has been defined, however, the distribution of polyyne biosynthetic gene clusters (BGCs), and their evolutionary relationship to alkyne biosynthesis, have not been addressed. We explored the distribution of alkyne biosynthesis gene cassettes throughout bacteria, revealing evidence of multiple horizontal gene transfer events. Investigating the evolutionary connection between alkyne and polyyne biosynthesis identified a monophyletic clade possessing a conserved seven-gene cassette for polyyne biosynthesis. Mapping the diversity of these conserved genes revealed a phylogenetic clade representing a polyyne BGC in *Pseudomonas, pgn*, and subsequent pathway mutagenesis and analytical chemistry characterised the associated metabolite, protegencin. In addition to unifying and expanding our knowledge of polyyne diversity, our results show that alkyne and polyyne biosynthetic gene clusters are promiscuous within bacteria. Systematic mapping of conserved biosynthetic genes across bacterial genomic diversity has proven to be a successful method for discovering natural products.

## Main

Bacteria and fungi are an unparalleled source of structurally and functionally diverse metabolites with important implications for medicine and agriculture. Different classes of metabolite can also possess common structural features, one such moiety is the presence of a single carbon-carbon triple (alkyne) bond. Many alkyne-containing natural products have been isolated from marine bacteria and possess biotechnology-exploitable spectra of biological activity [1]. Other metabolites possess elongated chains of alternating carbon-carbon single and triple bonds (polyynes). The sources of polyyne metabolites are highly diverse, and have been isolated from plants, fungi, and bacteria, with one report of a beetle-derived polyyne [2]. The first bacterial polyynes, cepacin A and B, were discovered in the bacterium *Burkholderia diffusa* (formerly *Pseudomonas cepacia)* [3]. However, the biosynthetic origin of cepacin was only defined recently in the closely related species *Burkholderia ambifaria,* and the metabolite shown to function in the biocontrol of damping off disease caused by oomycete *Globisporangium ultimum* [4]. The timeline of the discovery of bacterial polyynes is interesting, with multiple studies characterising molecular diversity and different ecological roles (**Fig. 1**). Following the discovery of cepacins A and B in 1984, several other polyynes have been elucidated from Proteobacteria. Caryoynencin was isolated from *Trinickia caryophylli* (formerly *Burkholderia caryophylli*) [5] and *Burkholderia gladioli* [6]. Alongside other anti-fungal compounds synthesised by *B. gladioli,* Lagriinae beetles exploit caryoynencin in a symbiotic relationship to protect their eggs from fungal attack [18]. Collimonins were discovered in *Collimonas fungivorans* and displayed antifungal activity [7, 8], and ergoynes were found in the marine grass endophyte *Gynuella sunshinyii* [9] **(Fig. 1**). The polyyne Sch 31828 isolated from the Actinobacteria phylum [10], and fischerellins A and B from the Cyanobacteria phylum [11, 12], both of their associated gene cluster remain unknown. While alkyne [13] and polyyne [6] biosynthetic mechanisms have been determined, the emergence of polyyne biosynthesis, their evolutionary relationship to alkyne biosynthesis, and overall polyyne diversity, have yet to be established.

**Figure 1.**
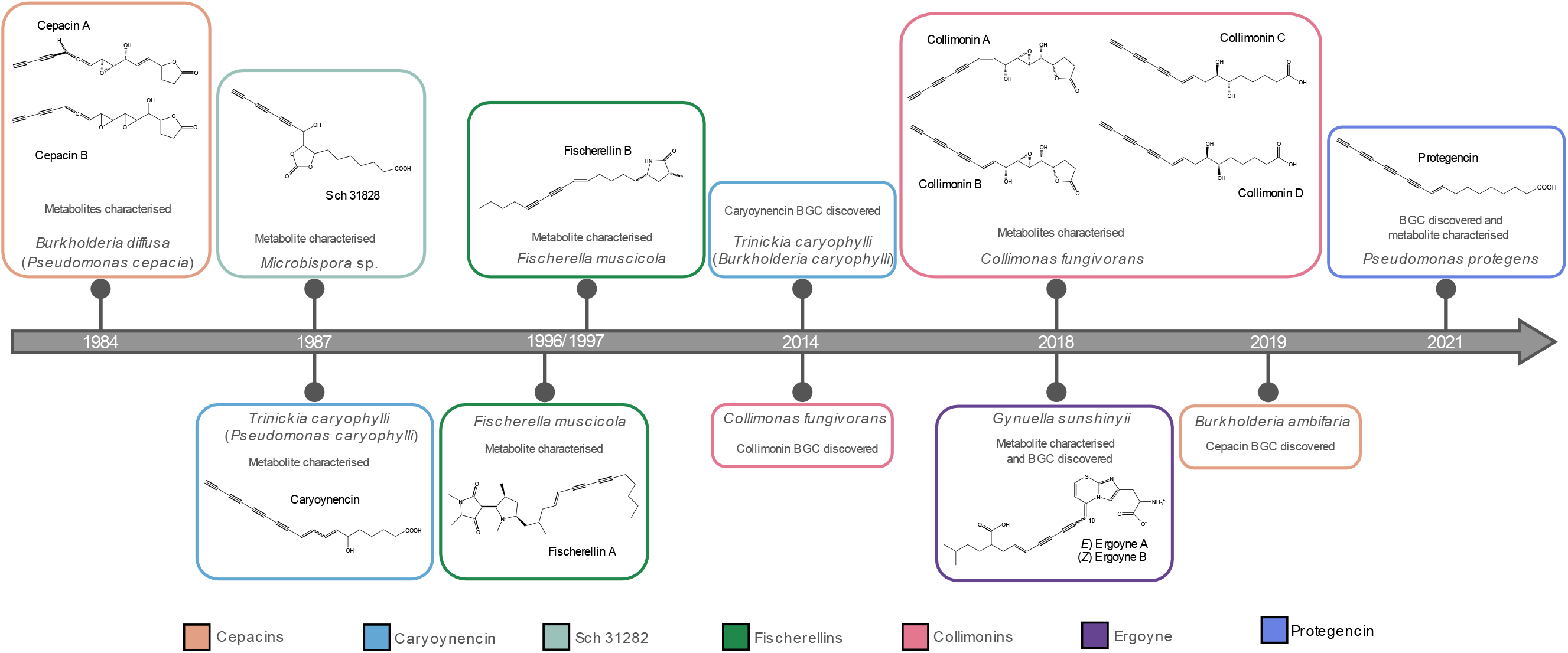
Timeline highlighting the discovery of polyyne metabolites and their biosynthetic gene clusters. The timeline displays the history of seven polyyne metabolites, their biosynthetic gene clusters, and the interval between the discovery of metabolite and BGC.

The influx of bacterial genomic assemblies over the last decade has revolutionised our understanding of bacterial evolution and enhanced our ability to discover natural products through multiple genome mining techniques (15). Following prediction of BGCs, common genomic approaches for characterising natural products include comparative metabolic profiling following mutagenesis of target BGCs, activation/inactivation of cluster-situated regulators, and heterologous expression (15, 16). Alternative methods fuelled by increasing availability of genomic data include analysing the evolutionary diversity of bacteria to identify lineages talented in specialised metabolite biosynthesis (15). A second, phylogeny-based mining strategy exploits the diversity of biosynthetic genes to discover natural product derivatives of known metabolites (15). Such an approach has the advantage of gleaming insight into the horizontal gene transfer of BGCs by comparing biosynthetic gene trees to evolutionary phylogenies.

Considering the limited study of polyyne evolution despite evidence of their evolutionary broad distribution (4, 11) (**Fig. 1**), we sought to integrate existing knowledge and expand our understanding of these structurally intriguing moieties. Here we show their evolutionary history by examining the co-occurrence of alkyne and polyyne biosynthetic cassettes, and their distribution, through a phylogeny-guided genome mining approach. Mixed evolutionary lineages within the alkyne phylogeny provided further evidence of their highly promiscuous nature. A distinct, monophyletic clade composed of polyyne biosynthetic gene clusters was observed within the broader alkyne gene cassette distribution. By examining sub-clade architecture, we identified a previously unexplored *Pseudomonas* polyyne clade that resulted in the characterisation of a novel polyyne BGC, *pgn,* and its associated metabolite, protegencin.

## Methods

### Detection of biosynthetic gene clusters, phylogenetic, and phylogenomic analyses

A BLASTp [20] search of NCBI genomes, excluding *Burkholderia* (taxid:32008) and a local database of *Burkholderia* assemblies (downloaded genomes and genomes assembled from publicly available Illumina read data) was performed with the cepacin homologue (CcnK [4]) of desaturase JamB as the query. The top 5,000 genus and species hits from NCBI were dereplicated, and their associated genomes downloaded and combined with the local collection. The flanking 30 kbp of protein hits with an E-value less than 1.00e^-50^ was extracted, and encoded protein domains predicted using Interproscan v5.38-76.0 [21]. Each sequence was screened for the presence of three domains corresponding to the presence of a fatty acyl-AMP ligase (IPR040097), fatty acid desaturase (IPR005804), and acyl carrier protein (IPR009081). The presence of these three homologues were considered evidence of alkyne biosynthesis potential. These sequence fragments were further screened for the presence of four additional protein homologues: two desaturases, a thioesterase, and a rubredoxin protein via BLASTp, to determine the potential for polyyne biosynthesis. Protein and nucleotide alignments were generated using MAFFT v7.455 [22], with the exception of core-gene alignments that were generated with Roary v3.13.0 [23]. FastTree v2.1.10 [24] and RAxML v8.2.12 [25] were used to construct phylogenies. Bacterial genomes were annotated with Prokka v1.14.5 [26]. Average nucleotide identity analyses (ANI) were initially performed with fastANI v1.2 [27] and supported by PyANI v0.2.9 [28]. A comparison of the annotated sequences was visualised using Easyfig [29]. Further details are available in the Supplementary Information.

### Mutagenesis of polyyne biosynthetic cluster

A range of in-frame, gene replacement, and insertional inactivation mutants were constructed in *P. protegens* and *T. caryophylli* (**Table S1**) to link polyyne biosynthesis to gene clusters and cassette function as described in the Supplementary Information.

### Metabolite extraction and LC-MS analysis of *P. protegens* wild types and *ΔpgnD* mutants

*P. protegens* wild type strains (CHA0 and Pf-5) and mutants (CHA0Δ*pgnD* and Pf-5Δ*pgnD*) were grown in LB medium at 30 °C overnight, and then inoculated onto PEM agar plates. After incubation at 22 °C for 3 d, the medium in single plate was cut into small pieces after removing cells and extracted with 4 ml of ethyl acetate (EtOAc) for 2 h, followed by rotary evaporation and re-dissolving in 1 ml of 50 % acetonitrile in water. The crude extracts were then analysed by UHPLC-ESI-Q-TOF-MS after centrifugation to remove debris. UHPLC-ESI-Q-TOF-MS analysis were performed using a Dionex UltiMate 3000 UHPLC connected to a Zorbax Eclipse Plus C-18 column (100 × 2.1 mm, 1.8 μm) coupled to a Bruker Compact mass spectrometer. Mobile phases consisted of water and acetonitrile (MeCN), each supplemented with 0.1% formic acid. After 5 min of isocratic run at 5% MeCN, a gradient of 5% to 100% MeCN in 12 min was employed with flow rate 0.2 ml min^-1^, followed by keeping constant for 5 min and then returning to initial conditions within 3 min. The mass spectrometer was operated in positive-ion or negative-ion mode with a scan range of 50–3,000 *m/z.* Source conditions were: end-plate offset at −500 V, capillary at −4,500 V, nebulizer gas (N_2_) at 1.6 bar, dry gas (N_2_) at 81 min^-1^ and dry temperature at 180 °C. Ion transfer conditions were: ion funnel radio frequency (RF) at 200 Vpp, multiple RF at 200 Vpp, quadrupole low mass at 55 *m/z,* collision energy at 5.0 eV, collision RF at 600 Vpp, ion cooler RF at 50–350 Vpp, transfer time at 121 μs and pre-pulse storage time at 1 μs. Calibration was performed with 1 mM sodium formate through a loop injection of 15 μl at the start of each run. Additional LC-MS methods are described in the Supplementary Information.

### Preparative HPLC purification and structure elucidation by NMR spectroscopy

*P. protegens* Pf-5 metabolite production was scaled up by extracting from 53 PEM agar plates (from 1.5 L medium). After growth at 22 °C for 3 d, the medium was processed as described for LC-MS analyses. The purification was performed on an Agilent 1200 Series HPLC instrument equipped with a HP Agilent 1200 Diode Array detector and an Agilent Zorbax C18 column (100 × 21.1 mm, 5 μm), and the crude EtOAc extract was separated with an MeCN–H_2_O gradient (0 min, 5% MeCN; 5 min, 30% MeCN; 50 min, 30% MeCN; 80 min, 100% MeCN; 90 min, 100% MeCN) at a flow rate of 9 ml/min and monitoring absorbance at 260 nm UV, which afforded a putative polyyne metabolite (1.5 mg, *t*_R_ = 76.8 min). The structure of this compound was elucidated on the basis of NMR spectroscopic data analysis. Sample was dissolved in 0.6 ml of deuterated solvent in a Norell^®^ standard series™ 5 mm NMR tube, and 1D/2D spectra (^1^H, ^13^C, COSY, HSQC, and HMBC) were obtained at 500 MHz for ^1^H NMR and 125 MHz for ^13^C NMR on a Bruker Avance III™ HD 500 MHz spectrometer. Chemical shifts (δ) are given in ppm, and coupling constants (*J*) are given in hertz (Hz). Additional HPLC methods are described in the Supplementary Information.

## Results

### Distribution of alkyne biosynthesis and emergence of polyyne biosynthesis

Phylogenetic trees based on 4990 representatives of the alkyne biosynthetic cassette proteins: fatty acyl-AMP ligase JamA, fatty acid desaturase JamB, and acyl carrier protein JamC, were constructed to assess the distribution of alkyne biosynthesis in bacteria (**Fig. 2**). All three phylogenies possessed the same broad topological structure, with most variation occurring within the central deep branches (**Fig. 2**). Phylogenies were also constructed based on the *jamABC* nucleotide sequences, which exhibited the same topological pattern to the proteinbased phylogenies (**Fig. S1**).

**Figure 2.**
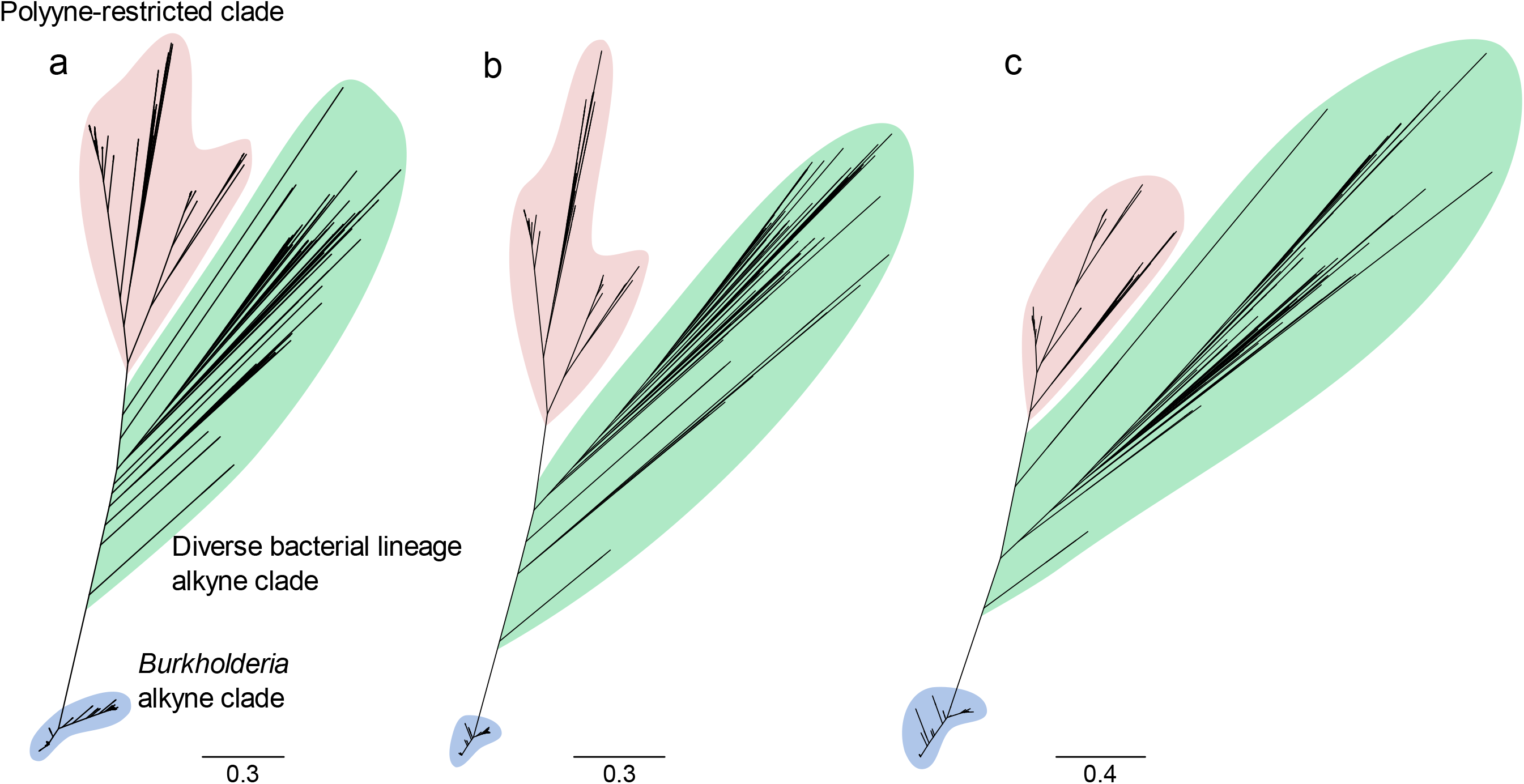
Protein-based phylogenies of potential alkyne-synthesising bacteria. Phylogenies were constructed based on 4990 representative of **a)** fatty acyl-AMP ligase JamA, **b)** desaturase JamB, and **c)** acyl carrier protein JamC homologues. The basal alkyne clade comprised of *Burkholderia* spp. is highlighted in blue, polyyne producers are highlighted in orange, and remaining deep-branching alkyne producers are highlighted in green. The general tree topology of the highlighted features is maintained across protein-based phylogenies, while the specific branching positions of sub-clades varies between phylogenies.

The ability to synthesise the alkyne moiety was widely distributed across Proteobacteria compared to other phyla, with representatives in most major phylogenetic clades potentially representing multiple acquisition events into the phylum (**Fig. 2** and **Fig. S2**). Within the Proteobacteria the alkyne biosynthesis cassette was predominantly found in Betaproteobacteria, but also included representatives of the classes Alpha-, Delta- and Gammaproteobacteria (**Fig. S2**). Outside of the Proteobacteria, examples of the cassette were found in members of the Cyanobacteria, Planctomycetes, and the candidate phylum Tectobacteria uncultivated sponge-symbiont “Entotheonella”. The phyla delimited clades suggest one main horizontal gene transfer event into each phylum. Most sequences (4013 sequences, approximately 80% of total) occurred within in a terminal clade composed entirely of *Burkholderia* species including *B. pseudomallei*, *B. thailandensis*, and *B. ubonensis* (**Fig. 2** and **Fig. S2**). A comparison of the JamA homologues extracted to those identified in the literature (14) highlighted a discrepancy in the JamA homologue previously defined in the *B. pseudomallei* alkyne biosynthetic locus, which was resolved by phylogenetics (**Fig. S3**).

To understand the broader relationship between bacterial alkyne and polyyne biosynthesis, a comparison of characterised polyyne biosynthetic gene clusters was performed. Analysis of the gene content and architecture of four characterised polyyne BGCs: cepacins, collimonins, caryoynencin, and ergoynes, identified seven common genes (**Fig. 3**). In addition to the three genes of the alkyne biosynthetic cassette, *jamABC* [13] two additional fatty acid desaturases, a thioesterase, and rubredoxin-encoding genes were shared between the BGCs (**Fig. 3**). Using this knowledge of conserved genes, we screened the flanking DNA sequence of the alkyne *jamABC* cassettes for the presence of the remaining four genes. This revealed a monophyletic clade in the alkyne phylogenies (**Fig. 2** and **Fig. S2**) where the 779 corresponding genomes possessed the conserved polyyne gene cassette (**Fig. 3**), with a few exceptions. Three discrepancies were observed within the monophyletic polyyne clade: *B. gladioli* strain 3848s-5 and three Streptomyces strains appeared to lack the co-localised thioesterase and rubredoxin genes with the remaining polyyne core biosynthetic genes, but manual inspection of these genomes revealed the BGCs were split across two contigs. A subset of 10 Actinobacteria genomes appeared to have replaced the thioesterase- and rubredoxin-encoding genes with a gene encoding a cytochrome P450 protein. These 10 genomes represented three genera and were confined to a single sub-clade in the monophyletic polyyne clade. The final discrepancy included two representatives of the family Mycobacteriaceae that lacked the rubredoxin gene.

**Figure 3.**
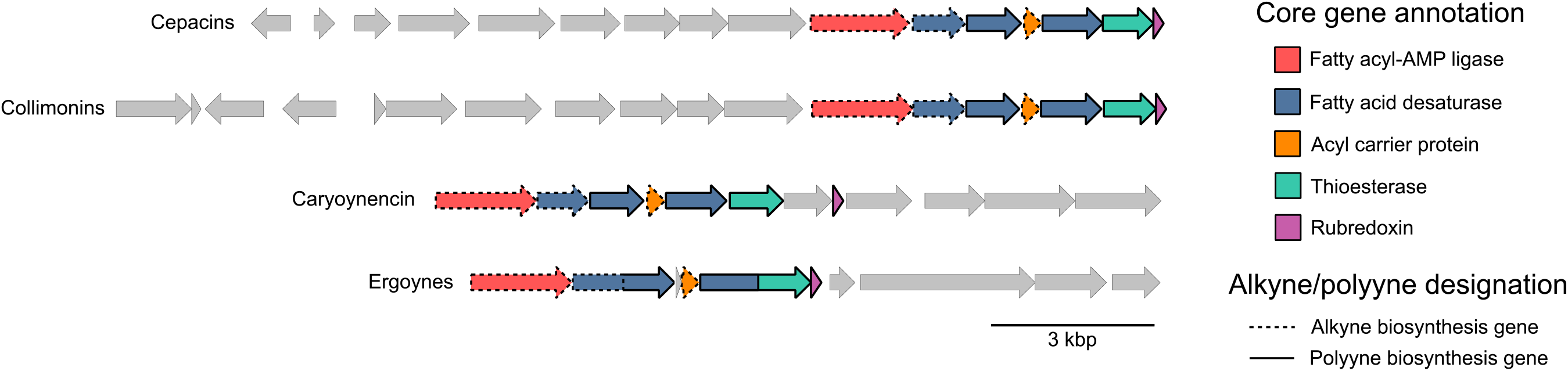
Comparison of gene organization between characterized polyyne biosynthetic gene clusters. Genes associated with alkyne biosynthesis: fatty acyl-AMP ligase (JamA), desaturase (JamB), and the acyl carrier protein (JamC) are indicated by a bold outline. Genes identified as polyyne biosynthesis specific genes: two desaturases and a thioesterase are indicated by a dashed outline.

To investigate the diversity of the monophyletic clade a separate phylogeny was constructed based on one of the polyyne-associated desaturase proteins (**Fig. 4**). This phylogeny was rooted using the basal branches of the clade of interest from both JamA and JamB phylogenies (**Fig. 2**): a Gammaproteobacteria sub-clade and Betaproteobacteria sub-clade. Within the resulting phylogeny we defined five major clades representing three Betaproteobacteria clades, one Gammaproteobacteria clade, and an Actinobacteria clade (**Fig. 4**). Each of the four previously characterised polyynes corresponded to a different clade, with collimonins, caryoynencin and cepacins localised to the three distinct Betaproteobacteria clades (**Fig. 4**). The ergoynes, synthesised by *G. sunshinyii,* were in the Gammaproteobacteria clade, but with deep-branching separating *G. sunshinyii* from the remainder of the clade members (**Fig. 4**). Each Proteobacteria clade was dominated by a single genus and mainly structured with relatively shallow branching. In comparison, the Actinobacteria clade possessed deep branching and contained representatives of seven genera including *Micromonospora*, *Actinomadura*, and *Rhodococcus,* but was dominated by *Streptomyces* species. This analysis identified the cepacin BGC in several species that were previously not known to carry the gene clusters (**Fig. 4**), and included *B. contaminans*, *B. vietnamiensis* and *Caballeronia peredens*.

**Figure 4.**
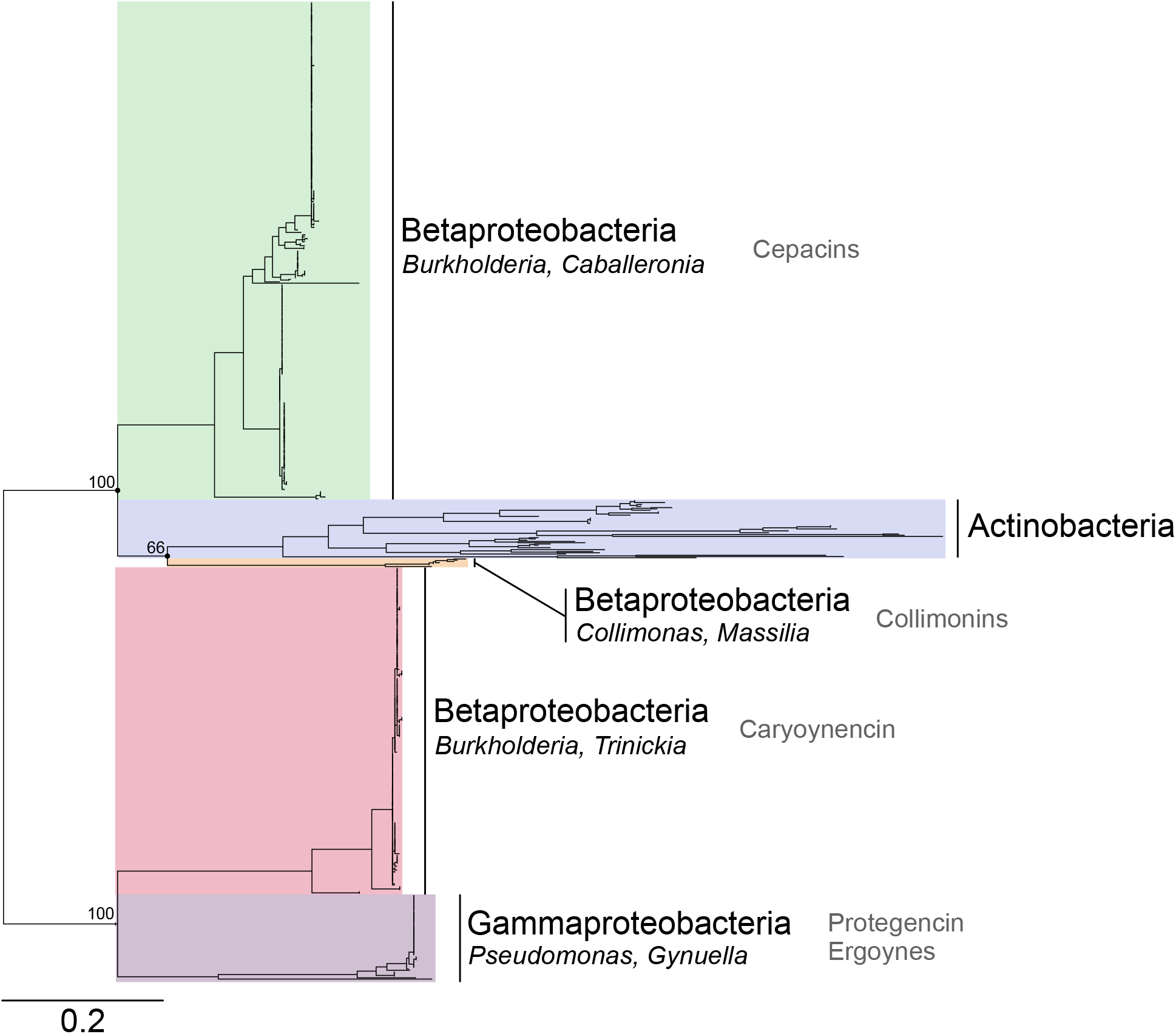
Desaturase protein-based phylogeny of polyyne producing bacteria. Homologues of the cepacin desaturase CnnN (protegencin PgnH) were extracted from bacterial genomes represented in the monophyletic alkyne clade as polyyne producers. The 4 Proteobacteria clades, their composite genera, and associated polyyne metabolites are indicated; in addition to the Actinobacteria phylum clade. The Gammaproteobacteria clade was used as the root based on the topologies of alkyne producer phylogenies using protein homologues of fatty acyl-AMP ligase (JamA), desaturase (JamB), and acyl carrier protein (JamC). Bootstrap values are indicated for splits between the 5 major clades. Scale bar represents substitutions per position.

### Exploration of the Gammaproteobacteria clade reveals an uncharacterised polyyne

Aside from the single representative of the *Gynuella* genus, the Gammaproteobacteria clade was dominated by *Pseudomonas*, however, no known polyyne has been characterised for this genus. Evidence of a *Pseudomonas* polyyne BGC has been alluded to as a homologous gene cluster of the collimonin [7] and caryoynencin [6] BGCs during the discovery of these polyynes. As such, we sought to investigate the potential of an uncharacterised bacterial polyyne in *Pseudomonas* (**Fig. 5a**), focussing on *Pseudomonas protegens* (formerly *P. fluorescens)* strains Pf-5 and CHA0 as our model bacteria (**Table S1**). HPLC analysis of these two strains revealed a small chromatographic peak with a characteristic UV-absorbance spectrum as observed in other polyynes [6, 7]. Mass ions identified via comparative negative ion mode HR-ESI-Q-TOF-MS analysis of the wild-type *P. protegens* Pf-5 and CHA0 strains against their respective unmarked deletion mutants of the fatty acyl-AMP ligase gene, Pf-5Δ*pgnD* and CHA0Δ*pgnD*, led to the identification of a compound which we named protegencin, with the molecular formula C18H18O2 (calculated for C_18_H_17_O_2_^-^: 265.1234, found: 265.1239) (**Fig. 5b-c** and **S4a-b**).

**Figure 5.**
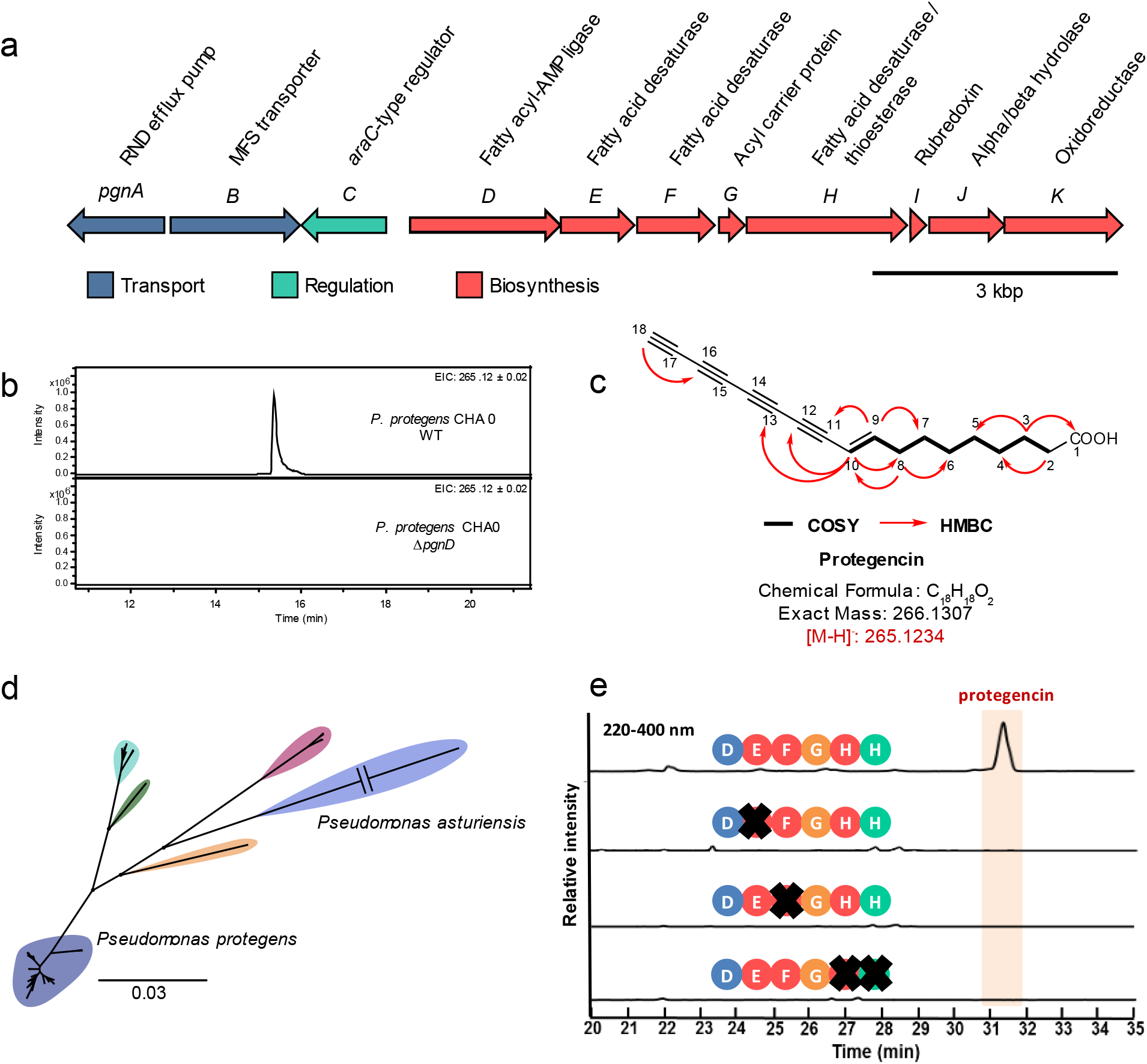
Organisation and distribution of the protegencin *(pgn)* BGC and analysis of protegencin production. **a)** Organisation and putative function of genes within the *pgn* BGC. **b)** Extracted ion chromatograms at *m/z =* 265.12 ± 0.02, corresponding to [M - H]^-^ for protegencin, from LC-MS analyses of crude extracts made from agar-grown cultures of *P. protegens* CHA0 (top) and the *P. protegens* CHA0 *ΔpgnD* mutant (bottom). **c)** Structure of protegencin, the identity of which was confirmed by a combination of high resolution mass spectrometry and NMR spectroscopy (see **Table S2** and **Fig. S4a-4g**). **d)** Core gene-based phylogeny, using 1487 genes, of 67 *Pseudomonas* genomes carrying the *pgn* BGC. The main nodes are highlighted that demarcate the *Pseudomonas* species, and all possess bootstrap values of 100. Scale bar represents substitutions per site. The *P. asturiensis* branch was shortened (indicated by a break) as such the scale bar does not apply. **e)** HPLC profiles (220-400 nm) of *P. protegens* Pf-5 wild type and inframe insertional mutant cultures. Only in the presence of all three desaturase genes *(pgnE, pgnF,* and *pgnH)* protegencin is produced. No polyyne precursor can be detected in the mutant strains.

### NMR spectroscopy confirms protegencin is a novel *Pseudomonas* polyyne

Polyyne compounds are notorious for being unstable and difficult to isolate, with recent studies requiring derivatisation by click chemistry before spectroscopic analysis [6]. The isolation of protegencin required a carefully optimised procedure to allow spectroscopic characterisation of the compound without derivatisation. Purified fractions of protegencin were dried under vacuum for 2 – 3 hrs, with addition of small volumes of MeCN to promote the removal of water from the sample. Freeze drying of protegencin-containing fractions resulted in a polymerised brown oil and should be avoided when working with polyyne compounds. Using this procedure, protegencin was isolated as a brownish, amorphous powder. Its ^1^H, ^13^C, COSY, HSQC, and HMBC spectra were acquired in DMSO-*d*6 (**Table S2** and **Fig. S4c-g**). The ^1^H NMR spectroscopic data displayed two olefinic protons (δ_H_ 6.65, 1H, dt, *J* = 16.0, 6.5, H-9; δ_H_ 5.79, IH, d, *J* = 16.0 H-10), a methine proton (δ_H_ 4.06, 1H, H-18), and seven pairs of methylene protons. The ^13^C NMR and HSQC spectroscopic data (**Table S2**) indicated 18 carbons, including three methine carbons (δ_C_ 155.4, 107.3, and 74.7), seven methylene carbons (δ_C_ 34.1, 33.4, 28.9 × 2, 28.8, 28.0, and 24.9), one carbonyl carbon (δ_C_ 175.0.), and seven quaternary carbons. The above data suggested a similar polyyne structure to caryoynencin [5] but lacking a pair of olefinic protons and an oxymethine proton. The structure was further established by COSY and HMBC spectroscopic data analysis (**Fig. S4f-g**). The HMBC correlations of H-9/C-II, C-8, and C-7, along with the couplings of H-10/C-9, C-11, C-12, C-8, and C-13 confirmed a double bond located at C-9/C-10 next to the polyyne scaffold, as observed in caryoynencin. The double bond at C-7/C-8 and hydroxyl group at C-6 in caryoynencin were missing in protengencin, which was clarified by HMBC correlations from a methylene (H_2_-8) to two methine carbons (C-9 and C-10) and two methylene carbons (C-6 and C-7), and from a methylene (H_2_-4) to two methylene carbons (C-6 and C-5), as well as COSY couplings of H_2_-8/H-9 and H_2_-7. The other COSY correlations of H_2_-3/H_2_-4 and H_2_-2, and of H_2_-4/H_2_-5, together with HMBC correlations of H-2/C-1, C-3, and C-4, and of H-3/C-1, C-2, C-4, and C-5, confirmed the structure of the saturated region of this metabolite. Therefore, the structure of protegencin was elucidated as shown in **Fig. 5c** as a novel polyyne natural product.

### Distribution of protegencin (*pgn*) BGC within *Pseudomonas*

Following the discovery of the previously uncharacterised polyyne metabolite, protegencin, we sought to fully understand the species distribution of the *pgn* locus. The *Pseudomonas* branches of the Gammaproteobacteria clade represented 67 *Pseudomonas* genomes. Subsequent average nucleotide identity analysis (ANI) of these genomes indicated the presence of multiple species. Based on the established 95% species delineation threshold for ANI [27, 36] six species were identified, these included two named species, *Pseudomonas protegens* and *Pseudomonas asturiensis,* and four unnamed species. The relatedness of these species to one-another is highlighted in the core-gene-based phylogeny (**Fig. 5d**). *P. protegens* was the dominant species possessing the *pgn* BGC, representing approximately 75% of available *pgn*-encoding genomes. A wider search for genome representatives of these six species in the European Nucleotide Archive (ENA) revealed that all genomes available of these species possess the protegencin (*pgn*) BGC, except for *P. asturiensis*. Of the two available *P. asturiensis* genomes, only the type strain LMG 26898^T^ contained the *pgn* BGC but it was absent from *Pseudomonas* sp. 286 (98.9% ANI to LMG 26898). The *pgn* locus is present in five out of six *Pseudomonas* species examined in this study.

### A conserved desaturase triad is essential for polyyne formation

The high conservation of the three desaturase genes and the thioesterase gene across all orthologous polyyne BGCs is remarkable (**Fig. 3**). To identify their roles, we performed targeted gene replacements. Specifically, we individually replaced the desaturase and thioesterase genes with a kanamycin and apramycin resistance cassette in the *P. protegens pgn* and *T. caryophylli cay* BGCs, respectively (**Fig. 5e** and **Fig. S5**). Sequence analyses indicated that pairs of desaturase genes *(pgnE/cayB* and *pgnF/cayC)* would have similar functions. The deduced gene product of *pgnH* codes for a didomain enzyme with desaturase and thioesterase functions that corresponds to *cayE* and *cayF*, respectively. The metabolic profiles of the mutant strains were compared by HPLC (220–400 nm) with those of the wild type strains, with or without the empty pGL42a or pJET1.2/blunt vector used during mutagenesis (**Fig. 5e** and **Fig. S5**). Whereas *P. protegens* Pf-5 (with or without the empty vector) produces protegencin, in the Δ*pgnE-Kan^R^*, Δ*pgnF-Kan^R^*, and Δ*pgnH-Kan^R^* mutant strains no polyyne precursor could be identified (**Fig. 5e**). Deletions of the desaturase genes *cayB, cayC,* and *cayE,* and the thioesterase gene *cayF* in *T. caryophylli* abolished the production of caryoynencin. The wild type (with or without an empty vector) generates the 7*E*/*Z*-isomers of caryoynencin, but the mutant strains *(ΔcayB-Apr^R^, ΔcayC-Apr^R^, ΔcayE-Apr^R^*, and *ΔcayF-Apr^R^* neither produce polyynes nor pathway intermediates (**Fig. S5**). These data indicate that the three desaturases and the thioesterase synergise in the production of polyynes. Interestingly, the same multienzyme system that gives rise to a tetrayne in the protegencin and caryoynencin BGCs, appears to form three triple bonds in the collimonin pathway, and 2 triple bonds and an allene moiety in cepacin pathway (**Fig. 1**).

## Discussion

### Highly transmissible alkyne and polyyne cassettes

Our results identify a single point of emergence of polyyne biosynthesis within bacteria and demarcate its evolution from alkyne biosynthesis (**Fig. 2**). The basal positioning of Proteobacteria within the polyyne phylogeny hints at a potential origin of the biosynthetic ability (**Fig. 4**), followed by horizontal gene transfer into Actinobacteria and other Proteobacterial classes. Additionally, the occurrence of alkyne biosynthesis genes across diverse bacterial lineages was also indicative of multiple horizontal gene transfer events. Few other biosynthetic capabilities appear to occur across a spectrum of bacterial lineages. However, the *trans-AT* polyketide synthases (PKSs), similar to alkyne biosynthesis, seem to have migrated between Proteobacteria and Actinobacteria (28).

### Phylogeny-driven metabolite discovery

Mapping the diversity of polyyne biosynthetic gene clusters through functional gene and protein phylogenies permitted the discovery of an uncharacterised *Pseudomonas* polyyne BGC derivative. The use of function-based phylogenies has been exploited previously to gain insight into natural product diversity, such as ketosynthase (KS) and condensation (C) domains of non-ribosomal peptide synthetase (NRPS) and polyketide synthase (PKS) BGCs (29). Genome mining of genes known to encode enzymes that synthesise specific structural moieties also enable discoveries and comparisons to other structurally related metabolites. A novel glutarimide-class metabolite, gladiostatin, was discovered in *Burkholderia* following identification of a BGC possessing genes similar to those associated with the biosynthesis of *Streptomyces* glutarimide antibiotics (30).

### Evidence of uncharacterised polyyne in *P. protegens*

We identified and characterised a novel *Pseudomonas* polyyne metabolite produced by the widely studied *P. protegens* strains Pf-5 and CHA0 (**Table S1**). *P. protegens* (formerly *P. fluorescens*). Both strains have an extensive history of biopesticidal properties [39, 40], indicative of the array of potent antimicrobial natural products synthesised by this species such as the antifungal metabolites 2,4-DAPG and pyoluteorin [39, 40]. Previous basic homology analyses had highlighted the existence of a polyyne BGC in *P. protegens* with similarities to the caryoynencin [6] and collimonin [7] BGCs. However, homology only to the core biosynthetic region was defined in these studies [6, 7] (**Fig. 2**), and the metabolic product was not identified. Additionally, a transcriptomics analysis of the Gac global regulatory system highlighted a locus possessing similarities to those in *Burkholderia* [41], with a gene organisation and putative gene functions like those found in the cepacin BGC [4].

Overall, we sought to understand the evolution and diversity of polyyne biosynthesis following emergence from the alkyne biosynthetic gene precursor cassette. This study exploited functional gene phylogenetics, alongside evolutionary analyses to explore polyyne biosynthetic diversity. Further, bioinformatics analyses supported by molecular biology and analytical chemistry led to the discovery and elucidation of a *Pseudomonas* derived polyyne BGC, *pgn,* and metabolite, protegencin. The conserved multienzyme system was proven to be essential for polyyne formation in both protegencin and caryoynencin biosynthesis (**Fig. 5e**). Discovering polyynes and investigating their biosynthesis will support future endeavours to understand these biologically active metabolites.

## Acknowledgements

A.J.M, G.W, J.Z, E.M, G.L.C, J.A.H.M and T.R.C acknowledge funding from the Biotechnology and Biological Sciences Research Council (BBSRC) grant references BB/S007652/1 and BB/S008020/1. A.J.M. acknowledges funding by the BBSRC South West doctoral training partnership (BB/M009122/1). T.R.C acknowledges funding from the Medical Research Council award MR/L015080/1 which funded the Cloud Infrastructure for Microbial Bioinformatics (CLIMB) used for data analysis. G.L.C. was the recipient of a Wolfson Research Merit Award from the Royal Society (WM130033). The Dionex 3000RS/Bruker MaXis Impact instrument used in this work were purchased with a grant to G.L.C from the BBSRC (BB/K002341/1). M.J. is supported by a BBSRC Discovery Fellowship (BB/R012121/1). C.H. and H.J.K. acknowledge funding by the Deutsche Forschungsgemeinschaft (DFG, German Research Foundation) - SFB 1127/2, ChemBioSys – 239748522 and the Pakt für Forschung und Innovation. We thank Gail Preston for supplying the strain *P. protegens* Pf-5 and George O’Toole (Dartmouth College, Hannover, USA) for provision of the pMQ30 mutagenesis strains and constructs. We thank the School of Biosciences Genomics Research Hub at Cardiff University for genome sequencing services, and project student George Mears for laboratory technical assistance during his BSc (Hons) final year project.

## Author Contributions

*Conceptualisation:* AJM, GW, EM, HJK, CH; *Data curation:* AJM, GW, JZ, MJ, HJK; *Formal analysis:* AJM, GW, YP, JZ, MJ, HJK; *Funding acquisition:* EM, GLC, JAHM, TRC, CH; *Investigation*: AJM, GW, YP, JZ, CR, MJ, HJK; *Methodology*: AJM, GW, YP, JZ, MJ, EM, HJK; *Project administration:* AJM, GW, JZ, GLC, EM; *Resources:* EM, GLC, JAHM, CH; *Software:* AJM; *Supervision*: EM, GLC, CH; *Validation*: AJM, GW, YP, JZ, MJ, HJK; *Visualisation*: AJM, GW, JZ, MJ, HJK; *Writing - original draft:* AJM, GW, YP, JZ, MJ, EM, HJK *Writing - review & editing:* AJM, EM, GW, TRC, MJ, GLC, HJK, CH

