## Supplementary Information for "Exploration of polyyne biosynthetic gene cluster diversity in bacteria leads to the discovery of the *Pseudomonas* polyyne protegencin"

### Supplementary Methods

#### Detection of alkyne and polyyne biosynthetic gene clusters

Genomes were downloaded from the European Nucleotide Archive (ENA) using a script from enaBrowserTools (https://github.com/enasequence/enaBrowserTools). The local assemblies constructed from Illumina paired-end fastq data were assembled via Shovill v0.9.0 (https://github.com/tseemann/shovill). Upon conformation of all three domains (IPR040097, IPR005804, and IPR009081) hidden Markov models (HMMs) and BLASTp were used to identify and extract homologues of proteins JamA, JamB, and JamC. A threshold of 1.00e-100 was used to determine the presence of the additional desaturase proteins based on a noticeable change in e-value between protein presence or absence. Manual analysis of the sequence fragments for the presence or absence of alkyne and polyyne associated genes was necessary to define the thioesterase and rubredoxin thresholds due to an indistinct change in e-value and BGCs occurring near contig edges.

#### Phylogenetic and phylogenomic analyses

In cases where the protein or gene sequence of interest occurred as a fusion, the region of interest was extracted for use in the alignment. Alkyne-related phylogenies were constructed using multithreaded FastTree v2.1.10 [24], while remaining phylogenies were constructed using RAxML v8.2.12 [25] with a general time reversible model and gamma distribution supported by 100 bootstraps.

#### In-frame deletion of fatty acyl-AMP ligase encoding-gene, pgnD, in Pseudomonas protegens

In brief, two fragments of approximately 500 bp flanking the fatty acyl-AMP ligase encoding-gene, *pgnD* (**Fig. 5a**), were PCR-amplified (**Table S3**) using Q5 DNA polymerase (NEB). The suicide vector pMQ30 was linearised by *EcoRI*/*HindIII* restriction digest (ThermoFisher) and co-transformed with the two PCR fragments into *Saccharomyces cerevisiae* YPH500 using the lithium acetate transformation method [31]. Overlapping homologous 30 bp regions between the PCR fragments and linearised pMQ30 vector enabled yeast-mediated homologous recombination. The yeast transformation mixture was then grown on synthetic defined agar with 2% w/v D-glucose and 7.7 g/L complete supplement uracil dropout mixture (Formedium) for 3 – 4 d at 30 °C. Plasmid extraction was performed with Zymoprep Yeast Plasmid Miniprep I (Zymo Research), and transformed into *Escherichia coli* DH5α with selection on LB agar supplemented with 10 µg/ml gentamicin. Colonies were screened by PCR (**Table S3**) for the correct allelic exchange vector and confirmed by Sanger sequencing (Eurofins Genomics). The recombinant allelic exchange vector was introduced to *P. protegens* via tri-parental mating using the *E. coli* helper strain HB101 carrying pRK2013. Transconjugants were selected by growth on M9 medium supplemented with 2% (w/v) sodium citrate and 25 µg/ml gentamicin, and incubation for 36 – 48 h at 37 °C. Transconjugants were confirmed by PCR (**Table S3**) to detect recombination in either the upstream or downstream region. A second recombination event was triggered by sucrose counter-selection by growth on non-salt lysogeny broth with 15% (w/v) sucrose at 30 °C. Loss of the integrated allelic exchange vector was confirmed by loss of gentamicin resistance. PCR targeting the fatty acyl-AMP ligase encoding-gene, *pgnD* was used to distinguish between *P. protegens* wild-type and mutants, and later confirmed by Sanger sequencing (Eurofins Genomics).

Q5-based PCR (NEB) thermal cycler conditions were as follows: Initial denaturation at 98 °C for 30 s; 40x cycles of denaturation at 98 °C for 10 s, annealing at variable temperature (**Table S3**) for 30 s, and extension at 72 °C for 60 s; then a final extension at 72 °C for 2 min. *Pseudomonas* genomic DNA was extracted using the Maxwell 16® instrument and tissue DNA extraction kit (Promega, UK). *Taq*-based PCR (ThermoFisher) thermal cycler conditions were as follows: Initial denaturation at 95 °C for 5 min; 35x cycles of denaturation at 95 °C for 30 s, annealing at variable temperature (**Table S3**) for 30 s, and extension at 72 °C for 2 min; then a final extension at 72 °C for 10 min. Genomic DNA was extracted using 5% (w/v) Chelex 100 resin [32].

**Construction and analysis of *P. protegens* Pf-5 gene replacement mutants**

*P. protegens* Pf-5 was cultivated on Luria-Bertani (LB) agar at 30 °C for 2 d and on LB broth at 37 °C with orbital shaking (150 rpm) for 1 d. Genomic DNA from pure culture of *P. protegens* Pf-5 was acquired using MasterPure$^{TM}$ DNA purification kit. A targeted double-crossover strategy was chosen to insert the kanamycin resistance gene amplified from pGEM-Kan between the flank regions of each gene to inactivate *pgnE*, *pgnF*, and *pgnH*. The corresponding genes in the putative protegencin BGC were PCR-amplified from the genomic DNA of *P. protegens* Pf-5 and ligated with the kanamycin resistance gene. The combined gene fragment was inserted into the pGL42a vector by T4 DNA ligase. *E. coli* TOP 10 cells were transformed with the above vector through electrophoresis at 2,250V, and selected through LB plates supplemented with kanamycin (50 μg/ml). After inoculation of selected *E. coli* cells in LB media with kanamycin, the vector containing the combined gene fragment was purified by using Monarch® Plasmid Miniprep Kit from NEB. The vector was introduced into *P. protegens* Pf-5 by electroporation at 2,500 V, and the transformant was incubated on LB agar containing kanamycin at 30 °C. The successful integration of the kanamycin resistance cassette was verified with specific primers (**Table S4**). Verified colonies were inoculated into LB medium with kanamycin at 30 °C until the $OD_{600\ nm}$ value reached 4–5. Then, the bacterial cells were collected by centrifugation (5,000 rpm, 5 min) and washed with Tris-Acetate-Phosphate (TAP) medium twice. *P. protegens* Pf-5 cells were incubated in TAP medium with kanamycin at 30 °C with orbital shaking (120 rpm) for 1 d.

Analytical HPLC was performed on a Shimadzu Prominence HPLC system consisting of an autosampler, high-pressure pumps, column oven and PDA using a Macherey-Nagel C18 reverse phase column (Nucleosil 100, 5 μm, 125 × 4.6 mm, flow rate 1 ml/min). HPLC-grade $CH_3CN$ and deionized water with 0.1% trifluoroacetic acid were used as mobile phase for HPLC. The gradient elution was $CH_3CH/H_2O$ with 0.1% (v/v) TFA 0.5/99.5 to 100/0 for 30 min, $CH_3CN$ 100% for 10 min. Preparative HPLC was performed on a Gilson Abimed equipped with Binary Pump 321 and 156 UV/Vis detector (eluent: water with 0.1% (v/v) TFA, acetonitrile) using a Macherey-Nagel C18 reverse phase column (Nucleosil 100, 5 µm, 250 × 10 mm, flow rate 5 ml/min). LC-MS measurements were performed using a QExactive Orbitrap High Performance Benchtop LC-MS with an electrospray ion source and an Accela HPLC system (Thermo Fisher Scientific, Bremen). For MS/MS measurements an Exactive Orbitrap mass spectrometer with an electrospray ion source (Thermo Fisher Scientific, Bremen).

**Construction and analysis of *T. caryophylli* gene replacement mutants**

A targeted double crossover strategy was chosen to insert the apramycin resistant gene cassette from pIJ773 between the flank regions of each gene to inactivate the *cayB*, *cayC*, *cayE* and *cayF*. Each gene in the *cay* gene cluster of was PCR-amplified from the genomic DNA of *T. caryophylli* and ligated with the apramycin resistance gene. The combined gene fragment was inserted into the pJET1.2/blunt vector and introduced into XL1-Blue *E. coli* competent cells. After inoculating of *E. coli* cells in LB media with apramycin and ampicillin selection markers, the vector containing the combined gene fragment was purified using the Monarch® Plasmid Miniprep Kit from NEB. The vector was introduced into *T. caryophylli* by electroporation at 2,500 V, and the transformant was incubated on potato dextrose agar (PDA) plates containing apramycin and ampicillin at 30 °C. Successful integration of the apramycin resistance cassette was verified with specific primers (**Table S4**). The verified colonies were inoculated into potato dextrose broth with apramycin and ampicillin at 30 °C for 1 d. The culture was incubated on potato dextrose agar with apramycin and ampicillin selection for additional 4 d at 30 °C. Ethyl acetate was added into the small pieces cut from the agar plate and kept for 2 h. The ethyl acetate extract was filtered and concentrated under reduced pressure. The analytic methods were same as *P. protegens* Pf-5 mutation studies.

**Pea exudate medium (PEM) preparation**

An agar composed of pea (*Pisum sativum*) seed exudate was prepared as a growth condition representative of the nutrient availability surrounding a germinating seed. Approximately 100 g of Early Onward variety *P. sativum* seeds were rinsed three times with de-ionised water, and finally suspended in distilled water made up to 500 ml. The seeds were incubated in the dark with agitation (40 rpm on a rocking platform) for 2 d at 22 °C. The seed exudate was removed and filtered twice, initially with a grade GF/D glass microfibre filter, and finally a grade GF/A glass microfibre filter. The filtered seed exudate was combined with distilled water at a 1:1 ratio, and purified agar (Oxoid) added to make a 1.5% agar when autoclaved.

**Table S1. Strains and plasmids used in this study for mutagenesis.**

| Strains/Plasmids | Function | Source or Reference |
|---|---|---|
| **Strain** | | |
| ***Escherichia coli*** | | |
| DH5α | Tri-parental conjugation donor strain | Invitrogen |
| HB101 pRK2013 | Tri-parental conjugation helper strain | Figurski and Helinski (1979) [14] |
| *E. coli* TOP10 | Competent cells for bulk preparation of pJET1.2- $Apr^R$ | Invitrogen |
| *E. coli* XL1-Blue | Competent cells for bulk preparation of pJET1.2- $Apr^R$ | Invitrogen |
| ***Saccharomyces cerevisiae*** | | |
| YPH500 (ATCC 76626) | Homologous recombination of allelic exchange vector | Pahirulzaman *et al.* (2012) [15] |
| ***Pseudomonas protegens*** | | |
| CHA0 (DSM 19095$^T$) | Wild-type for mutagenesis | German Collection of Microorganisms and Cell Cultures (DSMZ) |
| Pf-5 (ATCC BAA-477) | Wild-type for mutagenesis | Howell and Stipanovic.(1979) [16] |
| CHA0Δ*pgnD* | Mutant with clean deletion of fatty acyl-AMP ligase, *pgnD* | This study |
| Pf-5Δ*pgnD* | Mutant with clean deletion of fatty acyl-AMP ligase, *pgnD* | This study |
| Pf-5Δ*pgnE*-$Kan^R$ | *pgnE* gene replacement mutant with $Kan^R$ cassette | This study |
| Pf-5Δ*pgnF*-$Kan^R$ | *pgnF* gene replacement mutant with $Kan^R$ cassette | This study |
| Pf-5Δ*pgnH*-$Kan^R$ | *pgnH* gene replacement mutant with $Kan^R$ cassette | This study |
| ***T. caryophylli*** | | |
| Wild type (DSM50341) | Wild-type for mutagenesis | German Collection of Microorganisms and Cell Cultures (DSMZ) |
| Δ*cayB*-$Apr^R$ | *cayB* gene replacement mutant with $Apr^R$ cassette | This study |
| Δ*cayC*-$Apr^R$ | *cayC* gene replacement mutant with $Apr^R$ cassette | This study |
| Δ*cayE*-$Apr^R$ | *cayE* gene replacement mutant with $Apr^R$ cassette | This study |
| Δ*cayF*-$Apr^R$ | *cayF* gene replacement mutant with $Apr^R$ cassette | This study |
| **Plasmids** | | |
| pMQ30 | Allelic exchange vector | Shanks *et al.* (2006) [17] |
| pMQ30_F1_F2 | Recombinant allelic exchange vector with *pgnD* flanking homology arms | This study |
| pGEM-Kan | Source of kanamycin resistance gene ($Kan^R$) | Ishida, Lincke and Hertweck (2012) [18] |
| PIJ773 | Source of apramycin resistance gene ($Apr^R$) | Gust *et al*. (2003) [19] |
| pJET1.2 | Introduce $Apr^R$ to target organism | Thermo Fisher Scientific |
| pGL42a | Introduce $Kan^R$ to target organism | Lackner, Moebius and Hertweck (2011) [20] |

**Table S2. $^1$H (500 MHz) and $^{13}$C (125 MHz) NMR spectroscopic data of protegencin.**

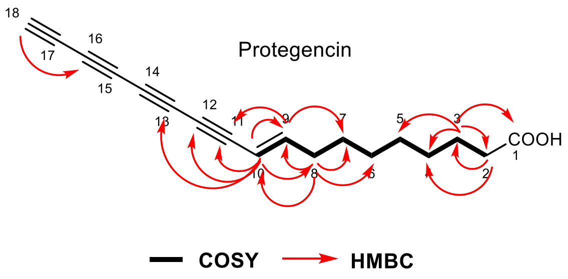


| Position | $\delta_H$ (*J* in Hz) | $\delta_C$, type |
|---|---|---|
| 1 | | 175.0, C |
| 2 | 2.18,overlapped | 34.1, $CH_2$ |
| 3 | 1.47, t-like (6.0, 6.5) | 24.9, $CH_2$ |
| 4 | 1.25, overlapped | 28.9, $CH_2$ |
| 5 | 1.35, overlapped | 28.8, $CH_2$ |
| 6 | 1.34, overlapped | 28.9, $CH_2$ |
| 7 | 1.35, overlapped | 28.0, $CH_2$ |
| 8 | 2.15, overlapped | 33.4, $CH_2$ |
| 9 | 6.65, dt (16.0, 6.5) | 155.4, CH |
| 10 | 5.79, d (16.0) | 107.3, CH |
| 11 | | 78.0, C |
| 12 | | 72.3 C |
| 13 | | 61.6, C |
| 14 | | 65.8*, C |
| 15 | | 60.3*, C |
| 16 | | 64.2, C |
| 17 | | 72.3*, C |
| 18 | 4.06, s | 74.7, CH |

*Might be interchanged.

**Table S3. PCR primers used for *P. protegens* clean deletion mutagenesis.**

| Primers | Final product size (bp) | Primer use | Ta (°C)[a]/DNA polymerase |
|---|---|---|---|
| **F1_fwd:** 5'-CACGACGTTGTAAAACGACGGCCAGTGCCAGGCCTGGACCTTGTCCAGTG-3'<br>**F1_rev:** 5'-TGGCCGTCATGACAGGTTCATTCAAAATCCTTTTTAAATGGAGAGCC-3' | 545 | Amplify upstream fragment (F1) | 62.5 / Q5 |
| **F2_fwd:** 5'-GGATTTTGAATGAACCTGTCATGACGGCCATGAATC-3'<br>**F2_rev:** 5'-ACACAGGAAACAGCTATGACCATGATTACGATCTGCATGATCAGTCGATC-3' | 550 | Amplify downstream fragment (F2) | 65 / Q5 |
| **M13_fwd:** 5'-AGGGTTTTCCCAGTCACGACGTT-3'<br>**M13_rev:** 5'-GAGCGGATAACAATTTCACACAGG-3' | 1,090 (pMQ30_F1_F2)<br>146 in (pMQ30) | Confirm recombinant pMQ30_F1_F2 construct | 58 / *Taq* |
| **M13_fwd:** 5'-AGGGTTTTCCCAGTCACGACGTT-3'<br>**Seq_chr_rev:** 5'-TACCGTCGGAATCGCCAGCC-3' | 3,132 (recombination at F1)<br>1,284 (recombination at F2) | Confirm transconjugant *P. protegens* | 60 / *Taq* |
| **Seq_chr_fwd:** 5'-CAGGACGAAAACTGCTGAACAGGA-3'<br>**M13_rev:** 5'-GAGCGGATAACAATTTCACACAGG-3' | 1,271 (recombination at F1)<br>3,119 (recombination at F2) | Confirm transconjugant *P. protegens* | 60 / *Taq* |
| **Seq_chr_fwd:** 5'-CAGGACGAAAACTGCTGAACAGGA-3'<br>**Seq_chr_rev:** 5'- TACCGTCGGAATCGCCAGC C-3' | 3,313 (wild-type)<br>1,465 (clean deletion) | Confirm *pgnD* clean deletion | 62 / *Taq* |

[a]Ta = Annealing temperature

**Table S4. PCR primers used for *P. protegens* and *T. caryophylli* gene replacement mutagenesis[a].**

| ***P. protegens*** | | |
|---|---|---|
| **Gene** | **Forward primer** | **Reverse primer** |
| *pgnE Fl1* | 5'- AAA AAA CTA GTG GCT GAC GGT CGC AAC TCC-3' | 5'- GCT ACT TAA TTA AGC TAG CGT AGC CCC TGG ATA TTG CCG ATA AA-3' |
| *pgnE Fl2* | 5'-GCT ACG CTA GCT TAA TTA AGT AGC ATA ACC CGC AGC TGG GAG-3' | 5'- AAA AAG AGC TCC AGA AGA ACT CGT CAA GAA GGC G-3' |
| *pgnF Fl1* | 5'- AAA AAA CTA GTA CAC AGC GAC ACC AGA GGT C-3' | 5'-GCT ACT TAA TTA AGC TAG CGT AGC ATC AGC ATC AGG CTG CTC ATC-3' |
| *pgnF Fl2* | 5'-GCT ACG CTA GCT TAA TTA AGT AGC ATC TTC GTC GCC AAC CAG GCG-3' | 5'- AAA AAG AGC TCT GCG GAC TTT GTC GAT CAT TG-3' |
| *pgnH Fl1* | 5'- AAA AAA CTA GTA CGC TCT CAG GAG ATC GAG GG-3' | 5'-GCT ACT TAA TTA AGC TAG CGT AGC TGC ATG TTG TGC ATG TGG C-3' |
| *pgnH Fl2* | 5'-GCT ACG CTA GCT TAA TTA AGT AGC AAC GGC TAC CTG AAG CTC-3' | 5'- AAA AAG AGC TCT TCG GCC ACC TTG ACG TTC TC -3' |
| *Kan[R]* | 5'-GCT ACG CTA GCG TAA GCT TAG GCT GCT GCC-3' | 5'-GCT ACT TAA TTA ATC AGA AA ACT CGT CAA GAA GGC G -3' |
| ***T. caryophylli*** | | |
| **Gene** | **Forward primer** | **Reverse primer** |
| *cayB Fl1* | 5'-ATC AGC ATC CGC ATG CGT C-3' | 5'-GCT ACT TAA TTA AGC TAG CGT AGC GTC GCC TTC CTG ATC GGT-3' |
| *cayB Fl2* | 5'-GCT ACG CTA GCT TAA TTA AGT AGC ATC GGC GGA ACT GCG TAC-3' | 5'-GCT CGA TGA ACG AGC CGG A-3' |
| *cayC Fl1* | 5'-ACG TAT CAA GGC GTT GAC CGC-3' | 5'-GCT ACT TAA TTA AGC TAG CGT AGC ACC CAG AAT AGA ATC GGA C-3' |
| *cayC Fl2* | 5'-GCT ACG CTA GCT TAA TTA AGT AGC ATT GCC AAT CAG TTC GCG-3' | 5'-GAC GCG GTC GAT CAT GT-3' |
| *cayE Fl1* | 5'-ATT CAG CTA CCC CGA CAG C-3' | 5'-GCT ACT TAA TTA AGC TAG CGT AGC ATG CAG GTT ATG CAT GTG-3' |
| *cayE Fl2* | 5'-GCT ACG CTA GCT TAA TTA AGT AGC ATC ACA TCT CGG ATG ATC-3' | 5'-CGA CTG GTA ATA GCC GTG CA-3' |
| *cayF Fl1* | 5'-ACA GAG CAG TCC ATT CTT CG-3' | 5'-GCT ACT TAA TTA AGC TAG CGT AGC AAC ATC GTG CTG CTG TCG-3' |
| *Apr[R]* | 5'-GCT ACG CTA GCA TTC CGG GGA TCC GTC GAC C-3' | 5'-GCT ACT TAA TTA ATG TAG GCT GGA GCT GCT TC-3' |

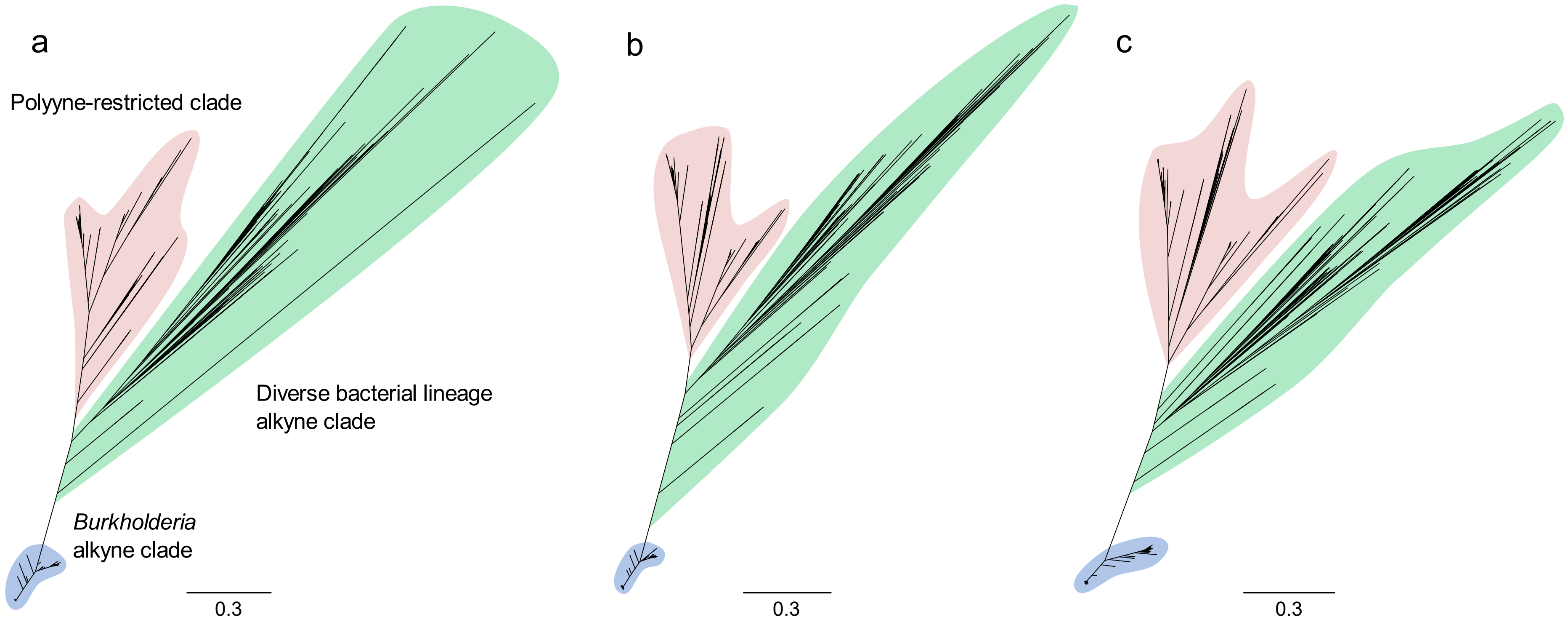


**Figure S1. Phylogenies based on alkyne associated genes.** Gene-based phylogenies were constructed based on **(a)** fatty acyl-AMP ligase gene *jamA*, **(b)** desaturase gene *jamB*, and **(c)** ACP gene *jamC* homologues. The basal alkyne clade comprised of *Burkholderia* spp. is highlighted in blue, polyyne producers are highlighted in green, and remaining deep-branching alkyne producers are highlighted in red. The general tree topology of the gene-based phylogenies reflects the protein-based phylogenies of JamABC.

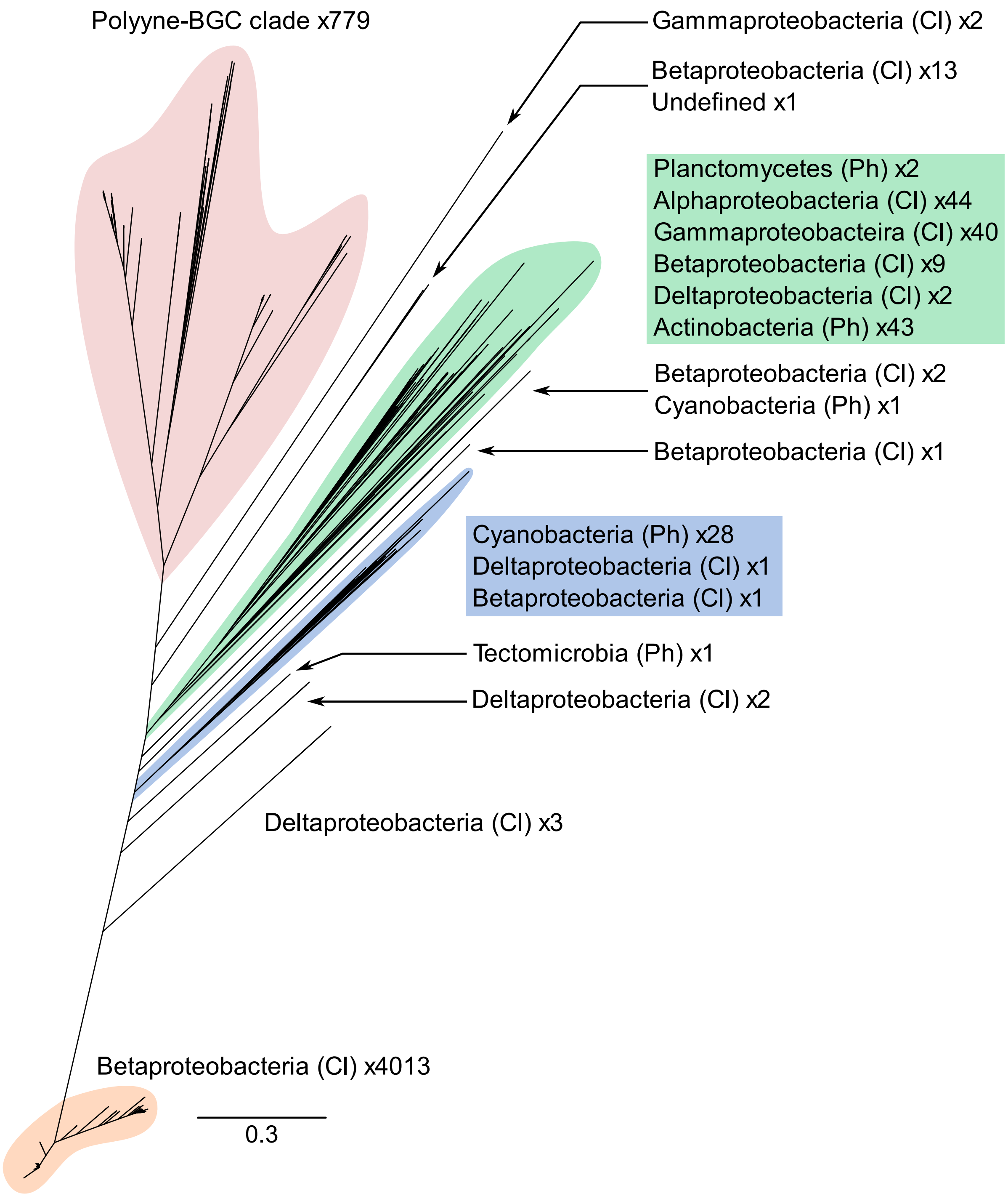


**Figure S2. Fatty acyl-Amp ligase (JamA) phylogeny of potential alkyne-synthesising bacteria.** The composition of each clade is indicated along with the number of representatives. Ph = Phylum; Cl = Class. Clades with high representation are highlighted with different colours. Same phylogeny as shown in **Fig. 2a**.

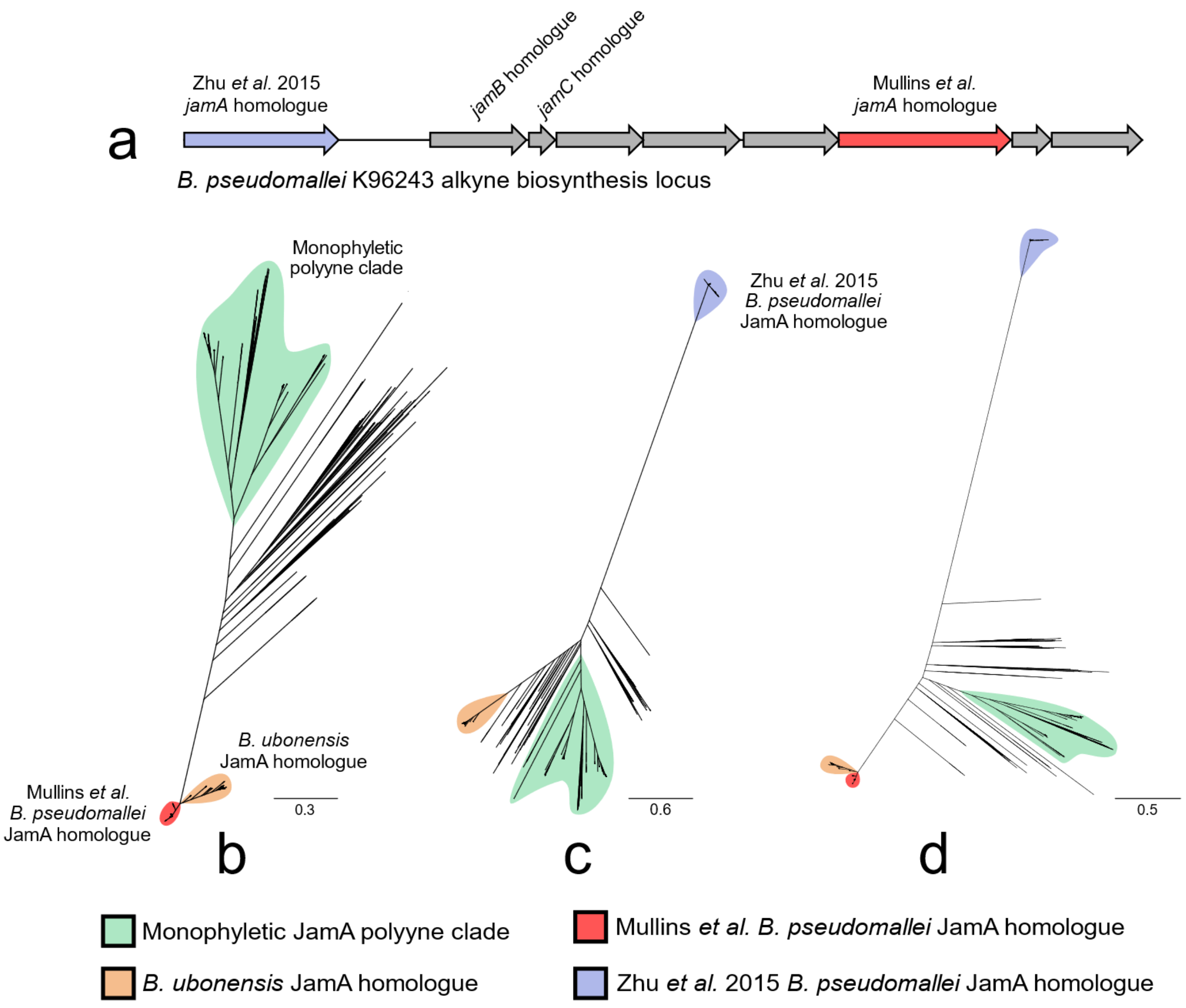


**Figure S3. FAAL protein-based phylogenies.** Comparison of the tree topologies of the JamA phylogeny with the presence and absence of the FAAL proteins identified in the *B. pseudomallei* alkyne biosynthesis locus. Key clades are indicated by colour to aid interpretation of tree topologies. **a)** Alkyne biosynthetic locus of *B. pseudomallei* K96243 with homologues of *jamABC* highlighted. **b)** Phylogeny of JamA homologues identified in this study (also displayed in **Fig. 2**). **c)** Replacement of *B. pseudomallei* JamA homologue identified in this study with the FAAL protein identified by Zhu *et al.* (2015) [21]. **d)** JamA phylogeny with *B. pseudomallei* FAAL proteins from this study and Zhu *et al.* (2015) [21] included.

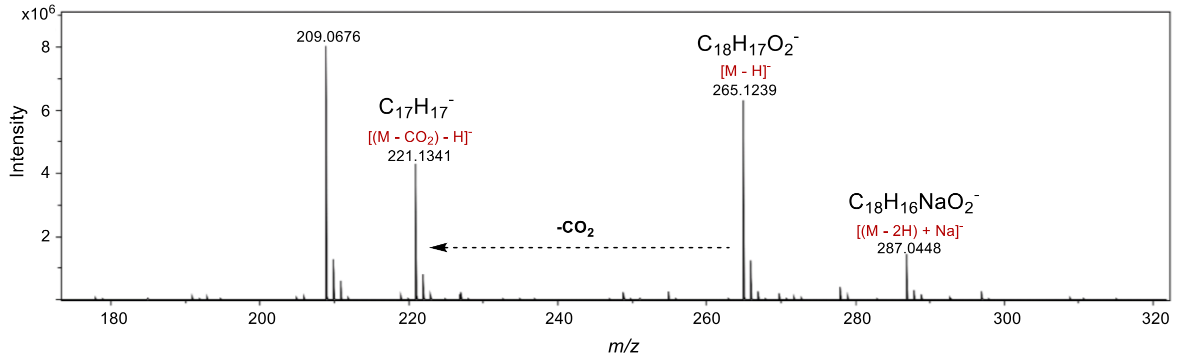


**Figure S4a: High-resolution mass spectrometry analysis of *P. protegens* CHA0 producing protegencin.** Measured spectrum of protegencin gives $[M\text{-}H]^-$, $[(M\text{-}2H)\text{-}Na]^-$ and $[(M\text{-}CO2)\text{-}H]^-$ions. The generated molecular formulae for each species are shown and are in agreement with the molecular formula of protegencin.

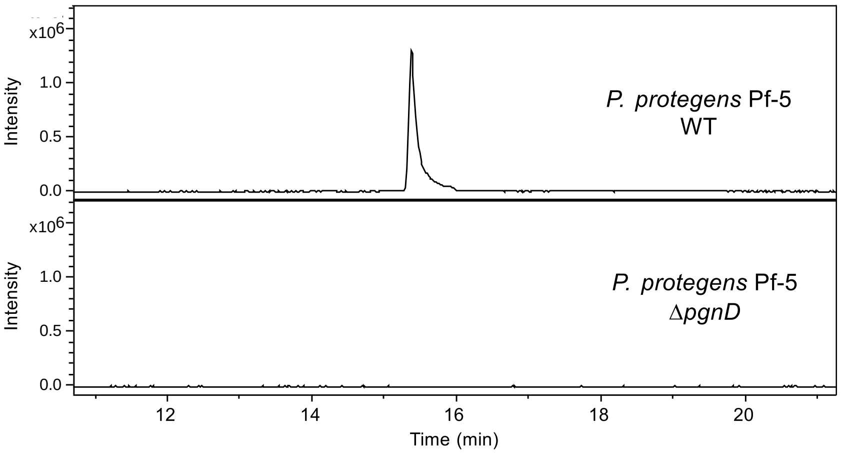


**Figure S4b: LC-MS analysis of protegencin production by *P. protegens* Pf-5.** Extracted ion chromatograms at *m/z*= 265.12 ± 0.02, corresponding to $[M - H]^-$ for protegencin, from LC–MS analyses of crude extracts made from agar-grown cultures of *P. protegens* Pf-5 WT (top) and the ΔpgnD mutant (bottom).

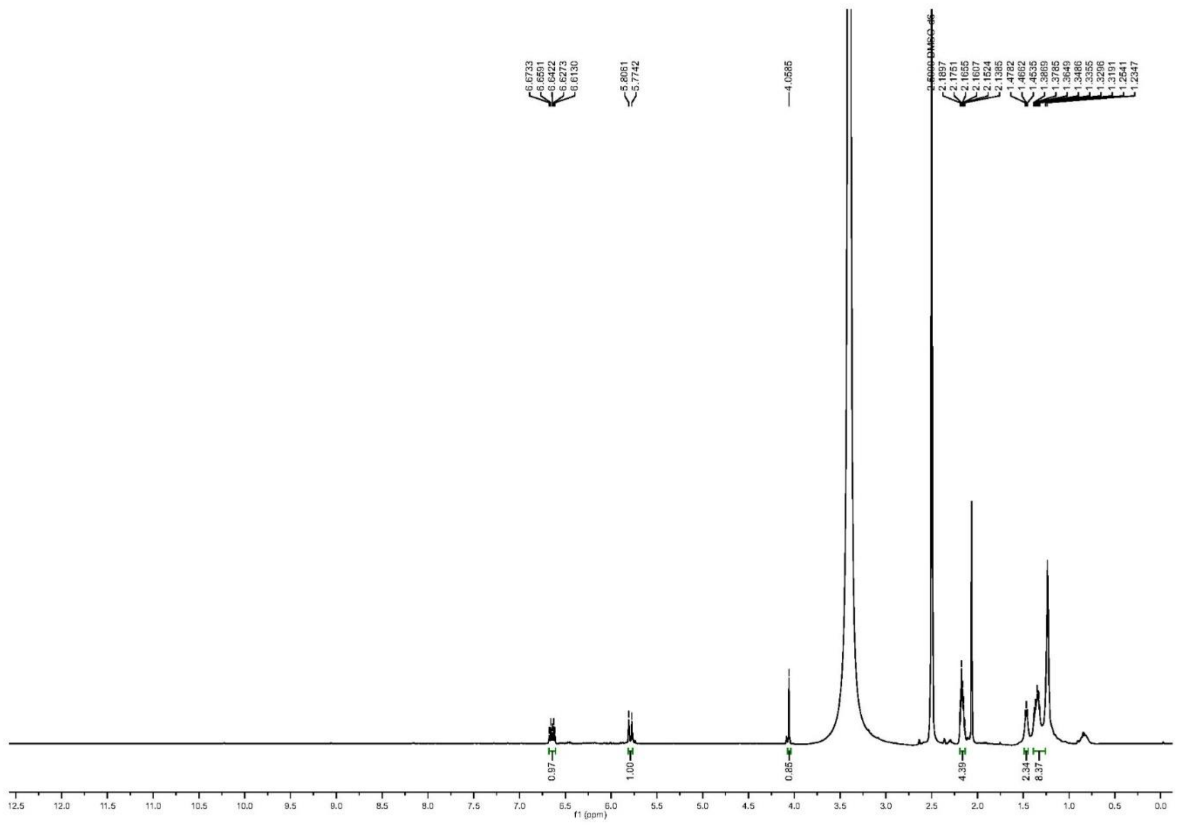

**Figure S4c: $^{1}$H NMR spectrum of protegencin in DMSO-d6 at 500 MHz.**

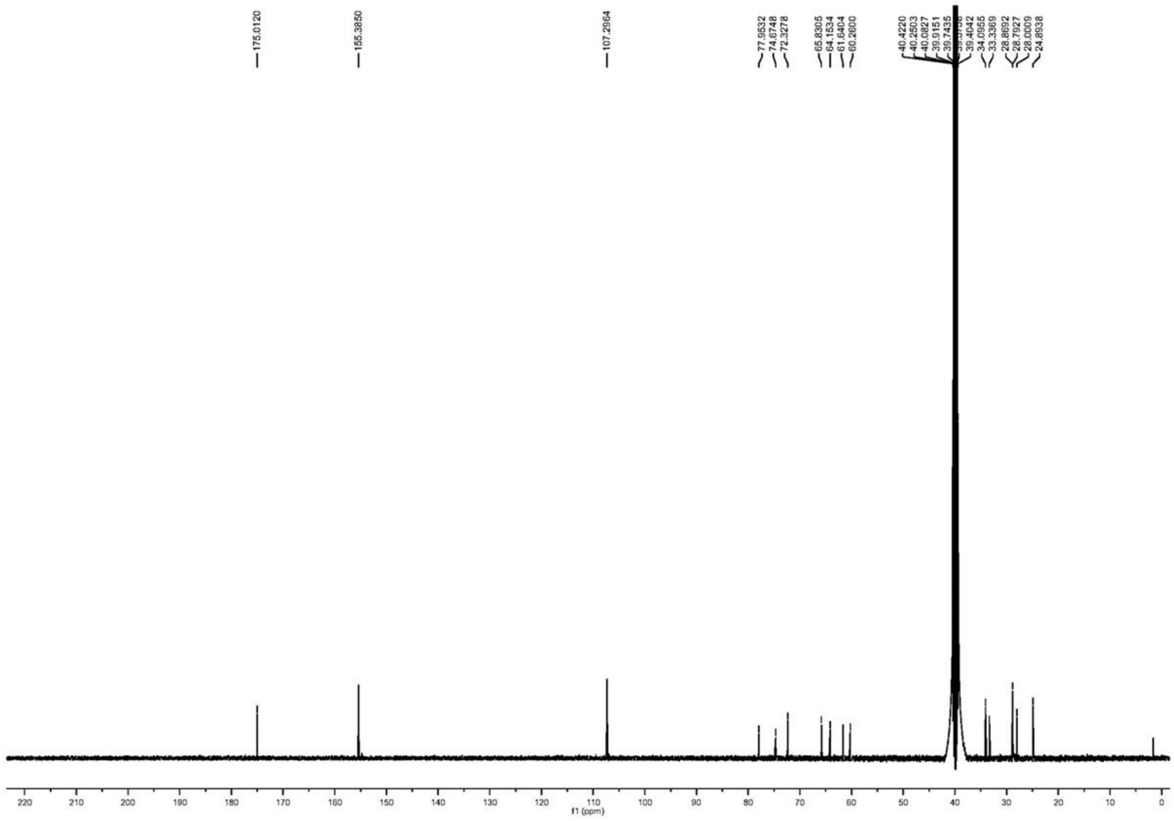


**Figure S4d: $^{13}$C NMR spectrum of protegencin in DMSO-d6 at 125 MHz**

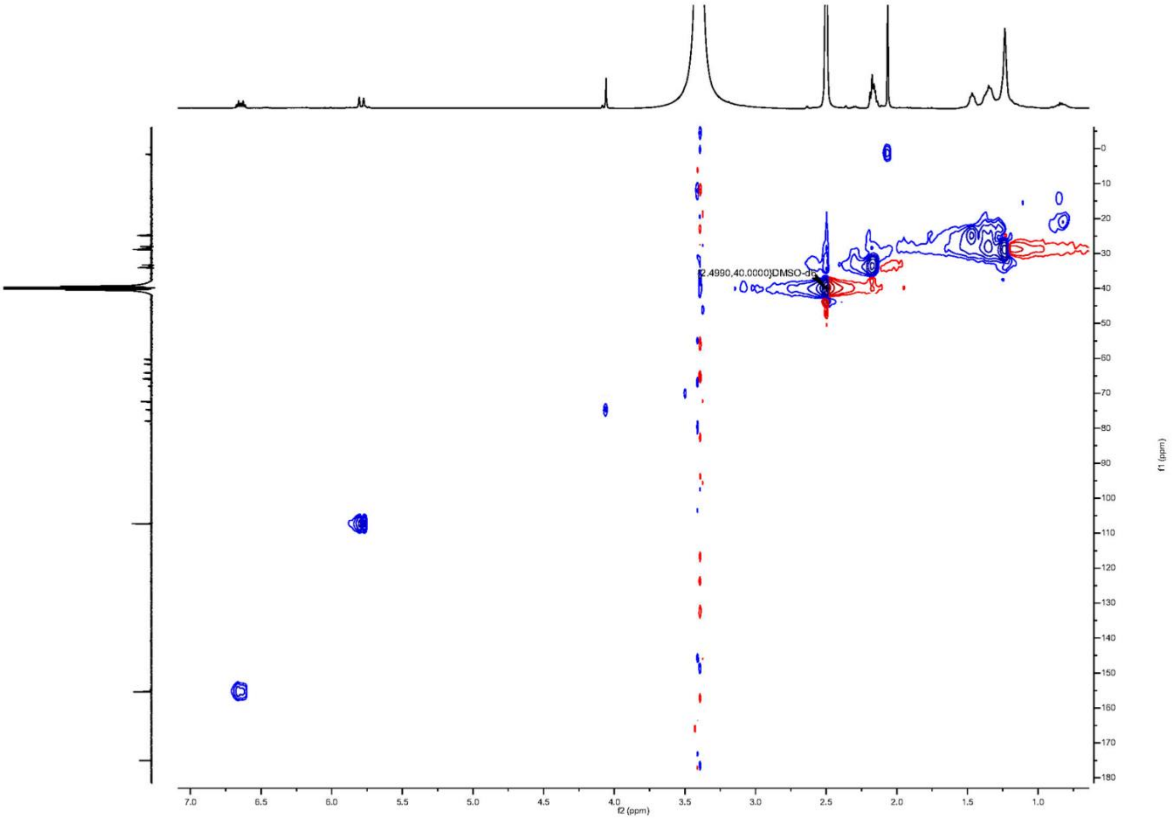


**Figure S4e: HSQC NMR spectrum of protegencin in DMSO-d6 at 500 MHz.**

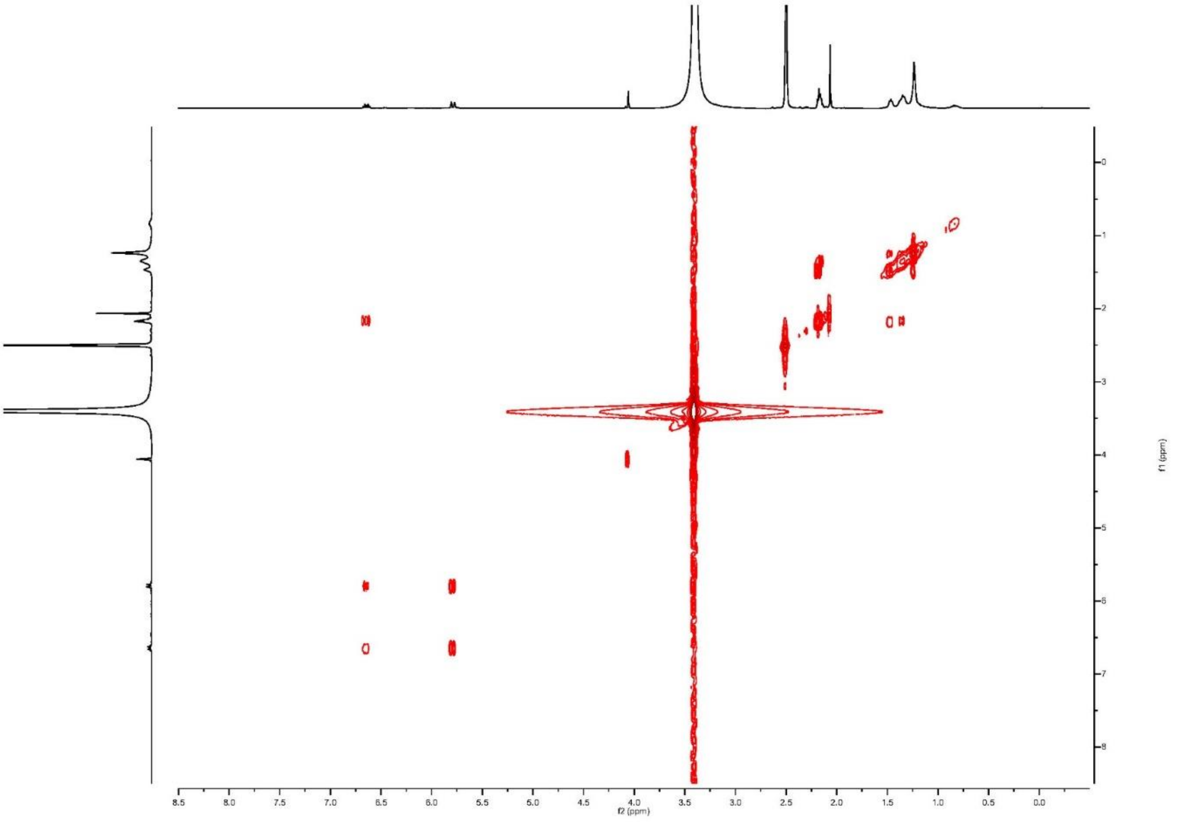


**Figure S4f: COSY NMR spectrum of protegencin in DMSO-d6 at 500 MHz.**

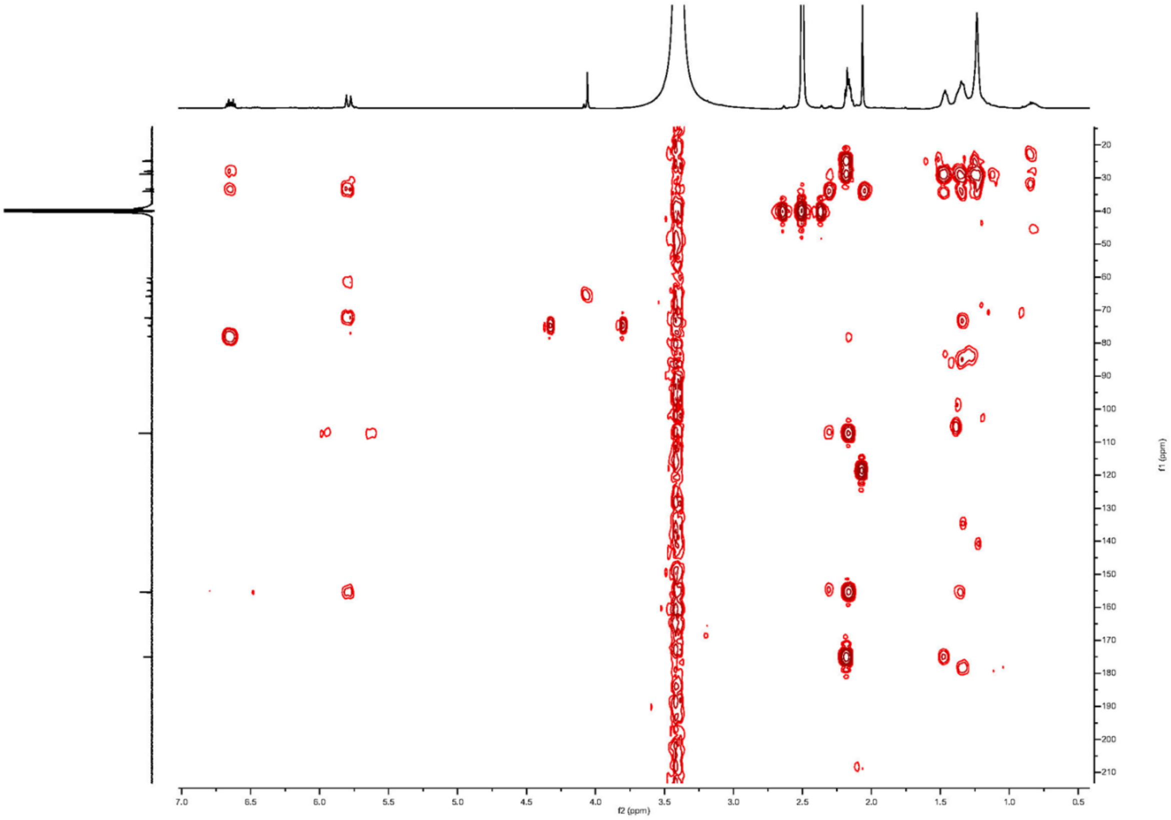


**Figure S4g: HMBC NMR spectrum of protegencin in DMSO-d6 at 500 MHz.**

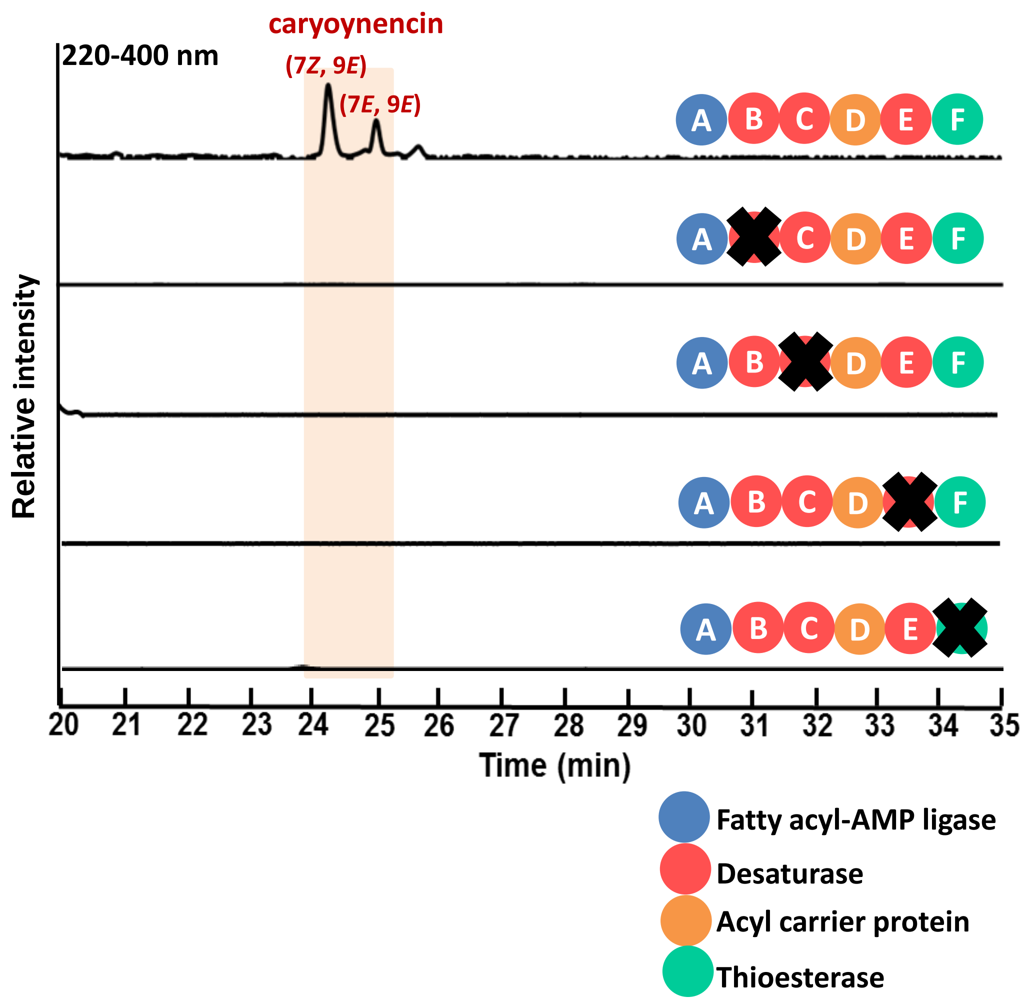


**Figure S5. HPLC analyses (220-400 nm) of the metabolite extracts from the *cay* mutants of *T. caryophylli.*** HPLC profiles of *T. caryophylli* wild type strain and mutants deficient in desaturase (*cayB*, *cayC*, and *cayE*) and thioesterase (*cayF*) genes. Caryoynencin can only be detected in the wild type culture. The mutant strain cultures (Δ*cayB*, Δ*cayC*, Δ*cayE*, and Δ*cayF*) do not produce any detectable polyyne precursors.
